# The demographic history of the wild crop relative *Brachypodium distachyon* is shaped by distinct past and present ecological niches

**DOI:** 10.1101/2023.06.01.543285

**Authors:** Nikolaos Minadakis, Hefin Williams, Robert Horvath, Danka Caković, Christoph Stritt, Michael Thieme, Yann Bourgeois, Anne C. Roulin

## Abstract

Closely related to economically important crops, the grass *Brachypodium distachyon* has been originally established as a pivotal species for grass genomics but more recently flourished as a model for developmental biology. Grasses encompass more than 10,000 species and cover more than 40% of the world land area from tropical to temperate regions. Given that grasses also supply about a fifth of the world’s dietary protein as cereal grains, unlocking the sources of phenotypic variation in *B. distachyon* is hence of prime interest in fundamental and applied research in agronomy, ecology and evolution. We present here the *B. distachyon* diversity panel, which encompasses 332 fully sequenced accessions covering the whole species distribution from Spain to Iraq. By combining population genetics, niche modeling and landscape genomics, we suggest that *B. distachyon* recolonized Europe and the Middle East following the last glacial maximum. Consequently, the species faced new environmental conditions which led to clear associations between bioclimatic variables and genetic factors as well as footprints of positive selection in the genome. Altogether, this genomic resource offers a powerful alternative to *Arabidopsis thaliana* to investigate the genetic bases of adaptation and phenotypic plasticity in plants and more specifically in monocots.

## Introduction

In the face of accelerating climate change, understanding how plant populations adapt to new environmental conditions has sparked great interest (Lasky et al. 2023). Environmental constraints varying in space and time alter the local frequencies of adapted genotypes, leaving detectable signatures in regions of selected genes. Identifying these signatures is crucial for understanding the processes underlying adaptation, and for providing candidate genes for functional studies. In this context, advances in genomics and DNA sequencing technology have revolutionized the field of landscape genomics by enabling high-resolution genome-wide analyses (for review Bourgeois and Warren 2021). For instance, genotype-environment association analyses (GEA) are powerful tools to identify alleles associated with ecologically relevant factors (Rellstab et al. 2015; Lasky et al. 2023). On the other hand, genome-wide scans of selection (Nielsen et al. 2005; Tang et al. 2007; Gautier 2015) have been largely used to identify genetic factors under selection without a prior neither on the phenotype under selection nor on the selective constraints acting on it. Those approaches, however, all rely on one fundamental aspect: the availability of genomic resources for many accessions occurring in contrasting habitats.

Despite the pressing need for new model species (Marks et al. 2023), research on the genetic bases and molecular characterization of local adaptation in plants is still dominated by *A. thaliana* (for review Provart et al. 2016; Woodward and Bartel 2018, Takou et al. 2019). The *A. thaliana* 1001 genomes project, combined with decades of functional studies, has indeed produced an unmatched wealth of knowledge about the processes of adaptation and evolution in plants (e.g. Hancock et al. 2011; Horton et al. 2012; Durvasula et al. 2017; Lee et al. 2017; Wu et al. 2017; Fulgione et al. 2018; Exposito-Alonso et al. 2019; Takou et al. 2019; Exposito-Alonso 2020; Wieters et al. 2021). Yet, grasses cover more the 40% of the land surface and play a key role in ecosystem functioning (Groves 2000). Developing alternative wild systems in monocots is thus very timely. Even though the rapid development of genomic resources for crops (Montenegro et al. 2017; Wang et al. 2018; Haberer et al. 2020; Jayakodi et al. 2020; Walkowiak et al. 2020; Lovell et al. 2021) will undoubtedly help to understand stress resilience in plants of agronomical interest, they nonetheless remain of limited value to tackle the diversity of paths to adaptation found in natural systems. In this context, we present here the *Brachypodium distachyon* diversity panel.

Initially established as a model for crop genomics (International Brachypodium Initiative 2010), the grass species *B. distachyon* is now a pivotal system for developmental biology (Hasterok et al. 2022; Raissig and Woods 2022) including the study of flowering time (Ream et al. 2014; Woods et al. 2014; Sharma et al. 2017; Woods et al. 2020; Bouché et al. 2022), stomata traits (McKown and Bergmann 2020; Nunes et al. 2020; Zhang et al. 2022; Slawinska et al. 2023) or cell-wall development (Coomey et al. 2020). In addition to harbouring a large mutant collection (Dalmais et al. 2013) and modern tools for mutagenesis (Hus et al. 2020), *B. distachyon* is a wild and widespread species with a small diploid genome (272 Mb). As such, it features functional and population genomics resources that are not combined in any other grass system for which a germplasm diversity panel has been established, such as rice or switchgrass (Wang et al. 2018; Lovell et al. 2021).

*B. distachyon* occurs naturally in oligotrophic habitats around the Mediterranean rim (López-Alvarez et al. 2015; Catalán et al. 2016). The combination of a geographic mosaic and climatic stability allowed Mediterranean species to diversify at regional and local scales (Nieto Feliner 2014). In this context *B. distachyon* constitutes an excellent system to investigate the genetic bases of local adaptation in a species adapted to arid climates. Earlier works on this topic mostly focused on a small set of accessions originating from Turkey and Spain (Del’Acqua et al. 2014; Des Marais et al. 2017; Bourgeois et al. 2018, Wyler et al. 2018; Stritt et al. 2018; Skalska et al. 2020) and the genetic basis of trait variation is still largely characterized through QTL mapping and mutant screening in this species (e.g. Barbieri et al. 2012; Woods et al. 2017; 2020; Jiang et al. 2017). Yet, genome-wide sequencing data have been produced for about 260 accessions originating from Spain, France, Italy, Turkey, Lesser Caucasus and Iraq (Gordon et al. 2017; 2020; Skalska et al. 2020; Stritt et al. 2022). Those resources, however, were never analyzed at once despite their great potential to unlock the source of genetic and phenotypic diversity in this species. In our previous study (Stritt et al. 2022), we analyzed a set of 196 accessions to provide a first insight into the population structure of this species. We showed that the expansion of three independent lineages during the Upper Pleistocene played an important role in the evolution of *B. distachyon* and further suggested that the interplay of high selfing and seed dispersal rates has shaped the genetic structure of this species. As a step further, we filled up here a last geographical gap by collecting and sequencing an additional set of *B. distachyon* accessions from Greece and Montenegro. We combined all available genomic data into a diversity panel encompassing 332 accessions and asked: i) What is the recent demographic history of *B. distachyon* populations ii) To what extent have environmental factors shaped genetic diversity in this species and, iii) Which genes have been selected by the environment. This resource is made available to the community to stimulate research and facilitate comparisons across established and emerging model species for plant adaptation.

## Results and Discussion

### Population structure and demographic history of *B. distachyon*

We used publicly available sequencing data from 196 natural *B. distachyon* accessions originating from Spain, France, Italy, Turkey, and Iraq (Gordon et al. 2017; Skalska et al. 2020; Stritt et al. 2022) previously analyzed in Stritt et al. (2022). We also added 65 additional sequencing data produced by Gordon et al. (2020) for accessions from Spain, Turkey and the Lesser Caucasus. Finally, we collected and sequenced an additional set of 71 accessions from Montenegro, Greece and France. Altogether, we assembled a diversity panel that encompasses 332 accessions (Fig. 1a, Table S1) for which whole-genome sequencing data are available with a minimum coverage of 20X.

**Figure 1.**
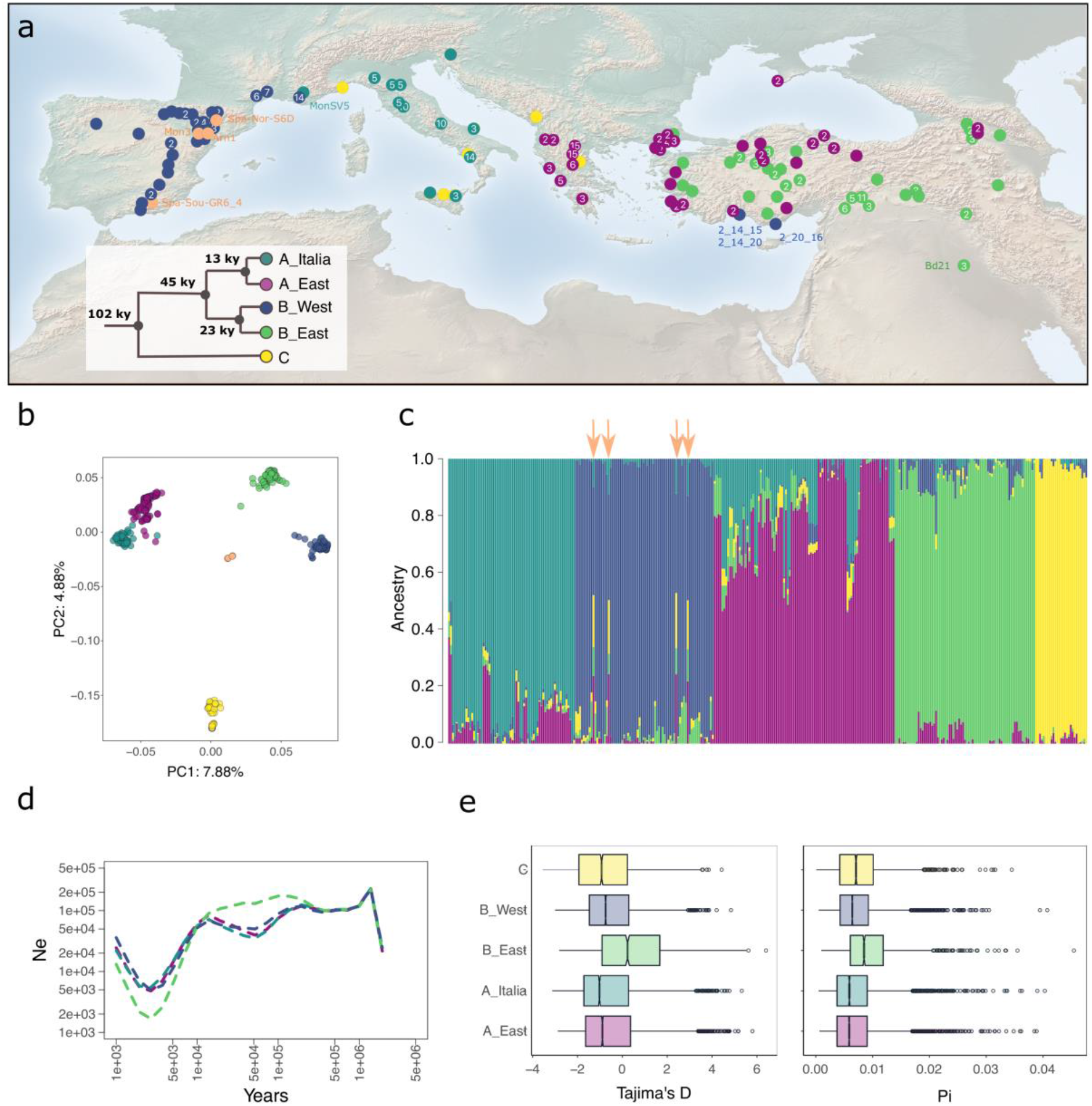
-Sample distribution and population structure. a) the map displays the origin of the 332 accessions used in the study. The number of samples collected at local sites is indicated in the circles. The tree represents the phylogeny of the five genetic clades and indicates the divergence estimates. The color code will apply to the rest of the study b) PCA based on 15,950 independent SNPs c) Inferred individual admixture coefficients. The black arrows point at the four admixed accessions found in Spain d) Population size evolution over time computed for the four derived genetic clades e) Tajima’s D and pi per genetic clade computed in 5kb windows over the entire genome.

*B. distachyon* belongs to the Brachypodium species complex. The three species in this complex, *B. distachyon, B. stacei* and the allopolyploid *B. hybridum*, can be difficult to distinguish morphologically but are straightforward to identify through genotyping (Giraldo et al. 2012; Catalán et al. 2016). Following extensive fieldwork, all accessions collected in the past in Morocco, Afghanistan, Iran, Pakistan, Australia and USA have been genotyped as *B. hybridum* (Wilson et al. 2019; Stritt et al. 2022; Fig. S1 for the geographical distribution of the 2,420 genotyped accessions). In contrast to *B. hybridum* and *B. stacei, B. distachyon* is extremely rare in Israel (Wilson et al. 2019; no Israeli samples included in the study) and in France only recorded in the South of the country (https://www.tela-botanica.org/bdtfx-nn-10075-synthese). The 332 *B. distachyon* accessions currently at hand (Fig. 1a, Table S1) are thus likely to cover the species range comprehensively.

We identified 10,227,760 high-confidence single nucleotide polymorphisms (SNPs) in the diversity panel and applied different filtering criteria according to the requirement of each analysis in the following sections, as described in Materials and Methods. While we merged datasets from different studies, a PCA based on pruned SNPs shows that samples do not cluster according to the study of origin (Fig. S2), indicating no large technical biases. Based on 196 samples, Stritt et al. (2022) found that *B. distachyon* accessions cluster in three main genetic lineages (A, B, C) that further split into five clades (A_East, A_Italia, B_East, B_West and C). Our current analysis based on 332 accessions did not reveal any additional discrete genetic clusters (Fig. 1a, b, c), even after expanding the sampling range. Indeed, the principal component analysis (PCA, Fig. 1b), sNMF (Fig. 1c and Fig. S3a,b) and the Evanno method (Evanno et al. 2005; Fig. S3c) all suggest that our accessions group into five main genetic clades.

Accessions from Spain and France share common ancestry and cluster together in the B_West clade (n = 72). One accession from France (MonSV5), however, shares common ancestry with the main Italian clade (A_Italia). Four accessions from Spain have a mixed ancestry and cannot be assigned to any of the clades (Spa-Sou-GR6_4, Spa-Nor-S6D, Mon3, Arn1). Accessions from eastern regions belong to two clades, one comprising Balkan and coastal Turkish individuals (A_East, n = 94), and the other including mainland Turkish and Anatolian individuals (B_East, n = 73). In Italy, two genetic clades, the A_Italia (n = 66) and C, co-occur. We also observed that all the accessions from Montenegro share ancestry with the rest of the C clade, as does one individual from Greece (Meg7; Fig. 1a-c). Altogether, the C clade comprises 27 accessions. Eleven accessions from the Lesser Caucasus have been identified as belonging to two different genetic clusters, with three accessions belonging to the A_East cluster and eight accessions belonging to the B_East cluster. Finally, three accessions from coastal Turkey (2_14_15, 2_14_20, 2_20_16) clustered with the B_West, as already reported in previous studies (Skalska et al. 2020; Stritt et al. 2022).

Using a multispecies coalescent approach, we estimated that the split between the ancestral C lineage and the A/B lineages occurred 102 thousand years ago (kya, 90% highest posterior density interval [HPDI] 50–170 ky), the split between the A and B lineages 45 kya (90% HPDI 21–76 ky) and the split within the A and B lineages 13 kya (90% HPDI 5-23 ky) and 23 kya (90% HPDI 11-39ky) respectively (Fig. 1a). These estimates are in the same order of magnitude as the ones we obtained with Relate, a method used for estimating genealogies genome-wide. For this analysis, the ancestral C clade was used to polarize SNPs. Relative cross-coalescence rates (CCR) between the remaining four genetic clades indicate that the split (CCR < 0.5) between the A and B lineages occurred within the last 100,000 years while the splits (CCR < 0.5) within the A and B lineages occurred within the last 5 ky (Fig. S4).

We also used Relate to compute effective population size (N_e_) evolution over time. Here again, the C clade was used for data polarization and therefore population size evolution could not be computed for this clade. This analysis revealed an abrupt decline in N_e_ across all clades around 30 kya, all followed by population expansion in a very recent past (Fig. 1d). Congruent with the recent population decline and expansion observed for each genetic clade, we found overall negative Tajima’s D values but still substantial remaining genetic diversity (pi) for all genetic clades (Fig. 1e). Nonetheless, the B_East clade displays higher Tajima’s D and genetic diversity levels than the other genetic clades (Kruskall-Wallis test, all p-values < 5.2e-05). On the one hand, Tajima’s D might remain slightly higher in the B_East clade because of the recent, more pronounced bottleneck. Gene flow/admixture are very limited in *B. distachyon* (Stritt et al. 2022) and unlikely to influence Tajima’s D. On the other hand, the slightly higher level of genetic diversity in the B_East clade could be explained by its more stable effective population size before the most recent drop/expansion ca. 5 kya. Note that all our dating methods rely on the use of a molecular clock and the assumption that plants reproduce only once a year. Although *B. distachyon* is annual, we have observed in the field that plants can set new flowers following grazing by sheep/goats. Hence, the assumptions made for these analyses might be approximated and our results must thus be interpreted with some caution. Even if the use of a strict molecular clock is arguable, the main results nonetheless indicate major changes in the recent demographic history of *B. distachyon*. Because these demographic changes accelerated within the last 10,000 years, we speculated that the species experienced a shift of its distribution following the Last Glacial Maximum (LGM, 22 kya) and more specifically during the Holocene period (11.7 kya) which marks the beginning of deglaciation in Europe.

### Ecological niche modeling and distribution of *B. distachyon* during the last glacial period

The climate, vegetation and landscape of Northern Eurasia (north of ca. 40°N and from 10°W to 180°E) underwent massive changes during the last glacial period (Binney et al. 2017; Davis et al. 2022). Under the cooler and dryer conditions faced during the LGM specifically, forests retreated to glacial refugia in Spain, Italy or the Balkans (for review Feliner 2011; Nieto Feliner 2014) and land remained mostly covered by steppe and tundra (Binney et al. 2017). Following the LGM, deglaciation led to the recolonization of Eurasia by woody plants and forests (Binney et al. 2017) and hence to substantial changes in dominant biomes. These recent climatic events have shaped the biogeography and genetic diversity of plants at large (Feliner 2011). In *A. thaliana*, for instance, relict populations occupied post-glacial Eurasia first and were later replaced by non-relict populations whose range expansions, accompanied by admixture, largely shaped modern populations (Lee et al. 2017).

To test for a potential shift in the distribution of *B. distachyon* in the recent past, we performed niche modeling analyses using Maxent climate niche models (Phillips et al. 2006). Due to the high correlation among bioclimatic variables for current time (Fig. S5a), we proceeded with a strategy based on selecting variables using an *a priori* understanding (Burnham and Anderson, 2004) of the *B. distachyon* life cycle: minimum temperature averaged from November to February (hereafter *tmin_Nov-Feb*) was chosen as potentially important for the vernalization process; precipitation levels averaged from March to June (hereafter *prec_March-June*), solar radiation levels averaged from March to June (*srad_March-June*) and elevation were chosen as relevant for plant phenology during the growing season.

We first fitted consecutive models for the whole species distribution with these four variables under current environmental conditions (see Materials and Methods), before projecting them over past conditions. Models with combinations of two variables had the lowest AIC values generally, however, only the model using *prec_March-June* and *tmin_Nov-Feb* showed Area under the Curve (AUC) values and predictive ability better than null models (p-value < 0.05) and was consequently chosen as our final model (see Fig. S5b for the distribution of these variables across genetic clades).

Based on this model and using paleoclimatic datasets, we projected the potential whole-species distribution in Europe, North Africa, and the Middle East since the Last Glacial Maximum (ca. 22 kya), as well as for the five genetic clades (Fig. S6). In contrast to the niche modeling performed with the geographically restricted current bioclimatic variables, we extended the analysis to a much wider geographic area in order to be able to detect putative changes in the species distribution. Our results suggest that climatic conditions similar to currently suitable conditions for *B. distachyon* extended southwards under LGM conditions with Northern Africa, Levant countries as well as the Iberic and Arabian Peninsula providing the most suitable habitats for the species (Fig. S6). During the LGM, the sea level was about 120–150 meters lower than at present (Yokoyama et al. 2000), which may have facilitated migration between Southern Europe and North Africa (Ortiz et al. 2007). Furthermore, Levant countries were mostly dry to subhumid habitats (Jennings et al. 2015). Hence, their geographical proximity with modern accessions and habitat suitability makes Northern Africa, Levant countries and the Iberic peninsula good candidates for *B. distachyon* glacial refugia. While the high suitability in the Arabian Peninsula might be seen as an overfit of our models at first, it is worth noting that annual rainfall levels under LGM conditions were much more important than nowadays in this region, making part of the Arabian Peninsula a dry to subhumid habitat (Jennings et al. 2015) likely suitable for *B. distachyon*.

We did not include information about soil as it is not projected for past conditions. For an oligotroph species as *B. distachyon*, the fundamental niche we computed might therefore differ from the realized one, and without modern samples available for all these four putative glacial refugia one can only speculate about a migration scenario (Lee et al. 2017). Yet, a natural conclusion is that *B. distachyon*, like most plants, operated a shift during the last glaciation and recolonized Europe and the Middle East following deglaciation. Our results contrast to some extent with those of López-Alvarez et al. (2015), who found that the distribution of *B. distachyon* extended southwards without a complete shift. This early study was based on the only samples collected at that time in Spain and Turkey. As it failed to recover the presence of the species in Italy and the Balkans under current environmental conditions, we argue that the projection to past conditions might be less reliable than the one we present here based on a more comprehensive sampling of the species.

The split among the three main genetic lineages of *B. distachyon* predates the LGM (Fig. 1a, Fig. S4) and it is therefore likely that accessions from the A_East/A_Italia, B_East/B_West and C genetic clades experienced different demographic histories. The geographic distribution of the five genetic clades is more parsimoniously explained by independent expansions and is sustained by the rapid increase of the effective population size for each clade within the last 5 ky (Fig. 1d). It has already been proposed that lineages and genetic clades may have used different migration corridors to recolonize Europe and the Middle East (Stritt et al. 2022). An East–West phylogeographical break, as often reported in Mediterranean species (Nieto Feliner 2014), could explain that accessions from the B_West and B_East genetic clades group together phylogenetically despite the current geographic gap (Fig. 1a). This scenario further supports a North African corridor for accessions from the B lineage, as already speculated in Stritt et al. (2022). Such a demographic scenario has been demonstrated in the herbaceous perennial species *Erophaca baetica* for instance, where plants from the Iberic peninsula are clearly derived from North African populations while geographically disconnected from the Greek and Turkish ones (Casimiro-Soriguer et al. 2010; Nieto Feliner 2014). In contrast, we previously observed a South-to-North gradient of declining N_e_, genetic diversity, and shared ancestry for A_Italia clade, which suggests a northwards expansion of this clade (Stritt et al. 2022).

### Genetic clades occupy different ecological niches and display adaptive loci

The Mediterranean basin comprises a mosaic of habitats (Nieto Feliner 2014) and consistent with recolonizations through different routes, the five genetic clades of *B. distachyon* occupy nowadays different geographical areas and ecological niches (Fig. 2). To detect significant differences among the ecological niche models (ENMs) of the five clades under current conditions, we created pseudoreplicate niche comparisons and compared them with the niche similarity score produced using the empirical data of the realized distribution (Fig. 2a-e). Note that the three accessions from Turkey clustering with the B_West clade (2_14_15, 2_14_20, 2_20_16, Fig. 1a) were excluded from the analysis. According to our models, there was no overlap between the niche suitability among the five clades. For the A_Italia vs C, B_West vs A_East, B_West vs A_Italia comparisons, p-values ranged from 0.01 and 0.05. For all the other comparisons, p-values were <0.01. We however acknowledge that the ENM of the C clade must be interpreted cautiously due to the small sample size.

**Figure 2.**
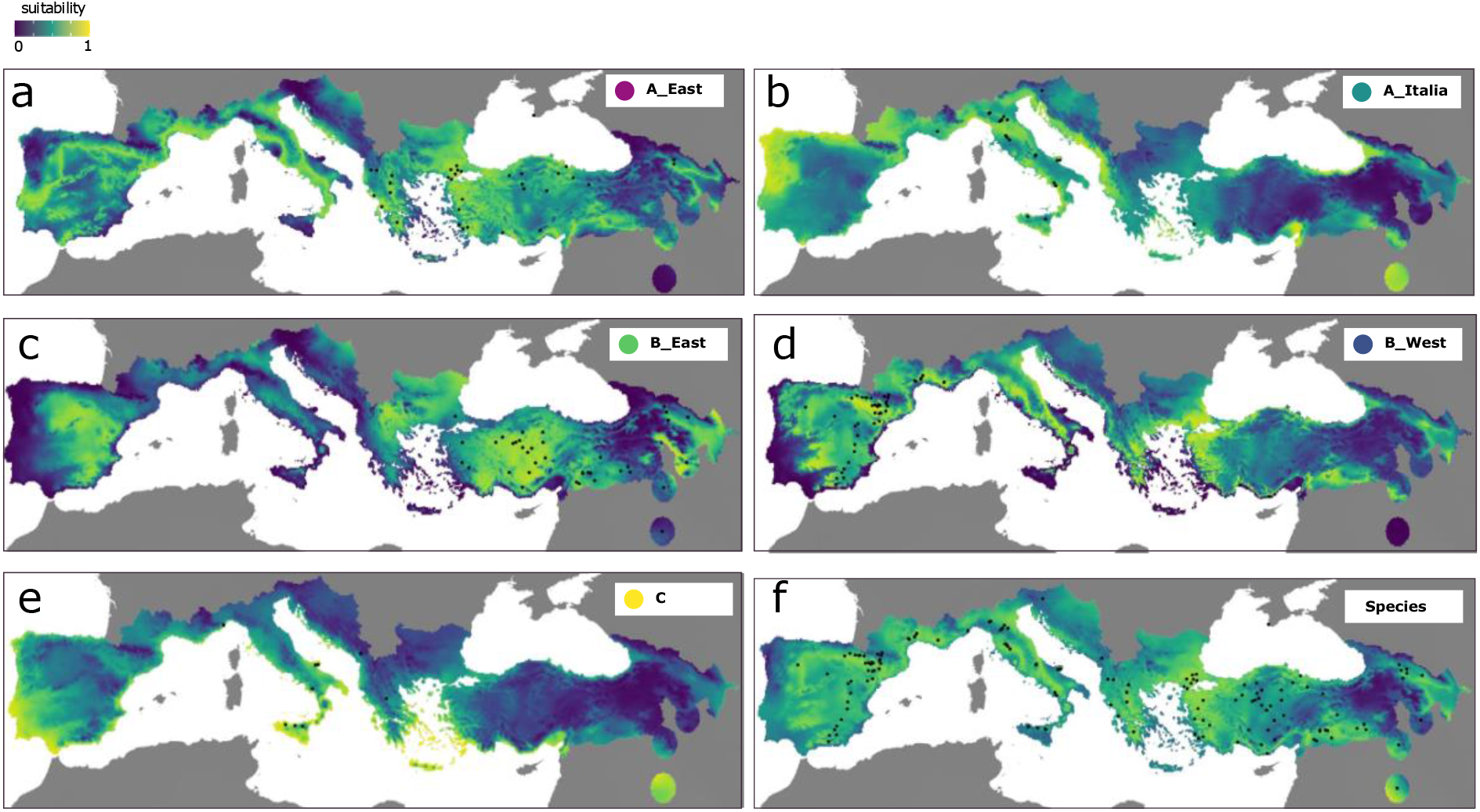
-Environmental Niche Modeling for current conditions using the masking region. The maps display environmental suitability for the five genetic clades a) A_East b) A_Italia c) B_East d) B_West e) C clade and f) at the species level.

More specifically, the higher predicted niche suitability for the A_East clade was at the coasts of Turkey, in mainland Greece, Italy, southern France, and parts of the Iberian Peninsula (Fig. 2a). The A_Italia clade displays a high suitability in Italy, Iraq, the northwester Iberian Peninsula as well as on the east coast of the Adriatic Sea (Fig. 2b). For the B_East clade, the areas with the highest suitability scores were mainland and eastern Turkey, northern Greece, coastal Bulgaria, as well as a part of central Spain (Fig. 2c), while the B_West clade displayed higher suitability in northern Spain, southern France, parts of Italy and Greece, as well as northwestern Turkey (Fig. 2d). Finally, the C clade shows high niche suitability in southern Italy, the southern Iberian Peninsula, the Greek Aegean islands, and Iraq (Fig. 2e).

We have already shown that adaptation at a regional scale led to specific footprints of positive selection in the genome of accessions from the B_East and B_West genetic clades (Bourgeois et al. 2018), but this previous analysis was somewhat limited by the number of samples then sequenced (27 and 17 accessions respectively). As we found that *B. distachyon* genetic clades now occupy different niches, we extended this initial study by testing for genome-wide footprints of positive selection at a regional level, using the genetic clades as focal populations. To do so, we calculated the XTX statistic, a measure comparable to single SNP Fst that accounts for the neutral genetic covariance across populations (Günther and Coop 2013) and detects highly differentiated alleles across several populations at once without SNP polarization into ancestral or derived alleles (Gautier 2015b). By selecting the top 0.1% XTX outliers (Fig. 3a), we identified 1477 highly differentiated genes putatively under positive selection (Fig. 3a and c). The GO annotation of the XTX gene set (Table S3) revealed a significant over-representation of genes involved in response to chemical (p-value = 2.01E-4) and metabolic process (p-value = 1.72E-04) including glutathione metabolic process (3.99E-06), a metabolic process involved in the control of reactive oxygen species (ROS) and hence stress response (Mittler et al. 2022).

**Figure 3.**
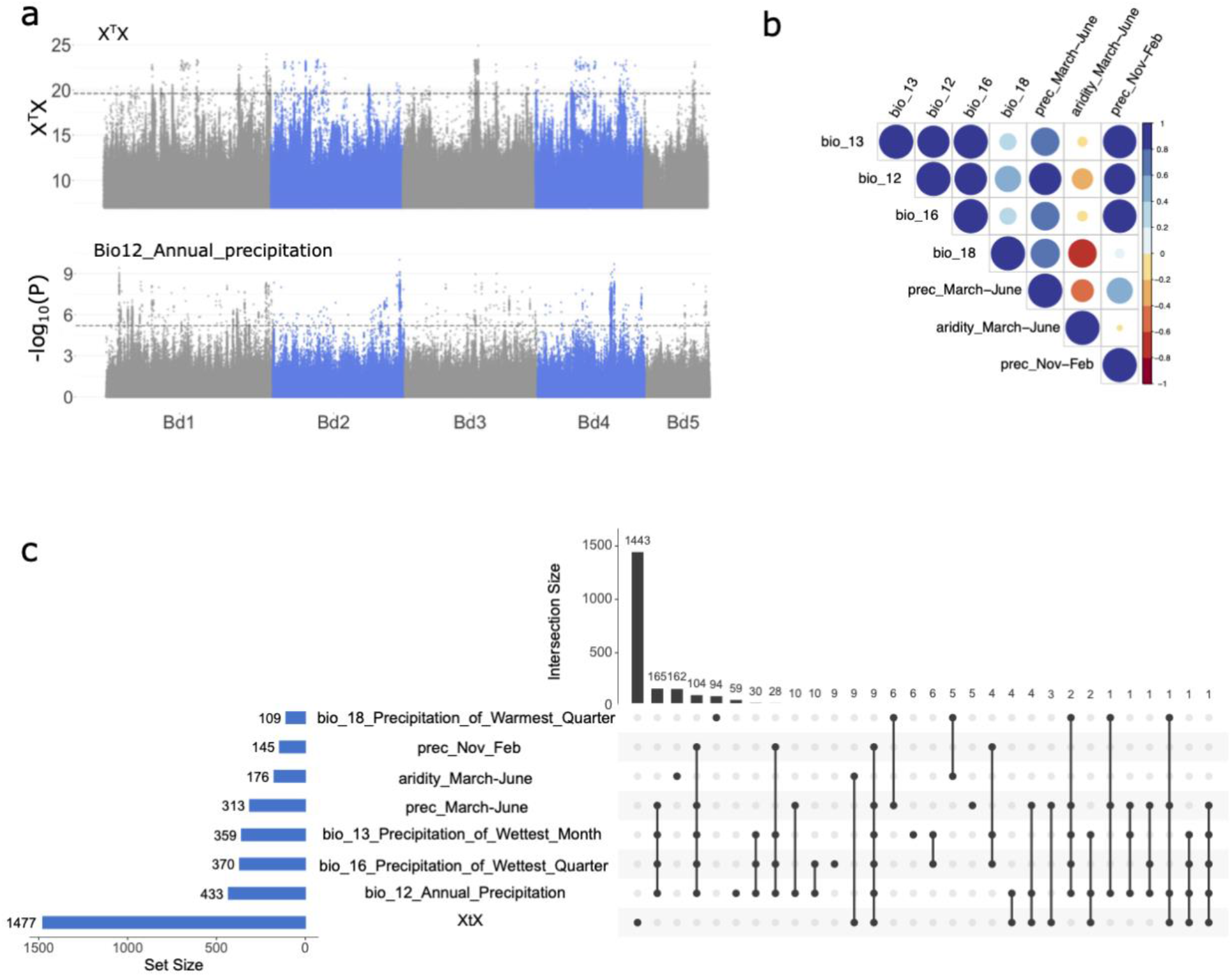
-Genotype-environment association and X^T^X analyses. a) Manhattan plots displaying regions under positive selection (the dotted line indicates the 0.1% outlier threshold) and the association between genomic regions and annual precipitation levels bio12 (the dotted line indicates the FDR threshold) b) Correlations among top 7 bioclimatic variables (associated with more than 100 genes) c) Upset plot displaying the overlaps among gene sets associated with bioclimatic variables or positive selection. The top 7 variables are presented. Set size shows the number of genes in significant regions of each specific variable. Intersection size shows the number of candidate genes associated with a variable (single dot) or shared among variables (multiple dots linked).

However, local adaptation is likely to occur at a much finer geographical scale (Gloss et al. 2022) and populations may each carry adaptations to their local climates (Lasky et al. 2023). Under the hypothesis that distinct lineages adapt to environmental gradients using the same traits (Lasky et al. 2023) through balancing selection, GEAs are powerful bottom-up tools to characterize the genetic basis of adaptation and identify which selective constraints might have shaped genetic diversity in a species. After excluding alleles displaying a minor allele frequency < 0.05, we tested for significant associations between the 2,867,335 remaining SNPs and 32 environmental variables related to precipitation levels, temperature, aridity index, solar radiation or elevation (Fig. 3b, Fig. S5a) with GEMMA while correcting for population structure (Zhou and Stephens 2012). We considered 5 kb windows significantly associated with a given environmental variable when they display at least two SNPs above the false discovery rate threshold (See Materials and Methods).

For each bioclimatic variable, we extracted genes located in significantly associated regions. Out of the 32,432 total genes already annotated in *B. distachyon* genome, 2,379 are significantly associated to at least one of the environmental variables (Fig. 3c, Fig. S7a, Table S2). Annual mean precipitation (bio12, Fig. 3a), together with bioclimatic variables associated to conditions from March to June (precipitation or aridity), yielded the highest number of genes (Fig. 3c). On the other hand, only 32 and 21 genes are located in regions associated with elevation or annual mean temperature (bio1) respectively. Interestingly, even though the 32 bioclimatic variables we chose show some levels of correlation (Fig. 3b, Fig. S5a), we only observed a partial overlap among the associated gene sets. For instance, while we identified 313 genes associated with precipitation levels in spring (*prec_March_June*) and 145 genes associated with precipitation levels in winter (*prec_Nov_Feb*), those two gene sets only share 113 genes indicating that genes playing a role in adaptation to climate may also be season-specific. The Gene Ontology (GO) annotation performed with the 2,379 genes associated with at least one bioclimatic variable (Table S3), however, did not reveal any significant process or molecular function.

We found little overlap between the GEAs and the scan of positive selection (Fig. 3c). This was expected given that we use the five genetic clades as focal populations for the XTX analysis and therefore putatively detect loci influencing adaptation at a larger geographical scale than with the GEAs. We also estimated the average age of alleles per gene with GEVA (Albers and McVean 2020). To avoid spurious age estimates, we only kept the 29,761 genes which harbor at least five SNPs (referred hereafter as genome-wide level). We used the gene lists produced by the GEA and the XTX analyses only if they contained at least fifteen genes. Even though statistical differences were observed among GEA and XTX gene sets, age estimates indicate that the large majority of alleles potentially involved in adaptation emerged 40 to 20 kya (Fig. S7b) and are slightly older than the recolonization of Europe and the Middle East by *B. distachyon*. In contrast to positive selection on *de novo* mutations (Barrett & Schluter, 2008), selection from standing variation is predicted to promote faster evolution (Hermisson and Pennings 2005) and this latter result suggests that the recycling of older alleles may have played an important role in local adaptation in our system.

### Current limitations and perspectives

We have not considered here that plants may also adapt along gradients via distinct strategies and traits, as shown in *A. thaliana*, where genetic factors causing local adaptation to elevation vary across regions and populations (Yan et al. 2021; Gamba et al. 2022). Because we filtered SNPs with more than two alleles, we may have generally excluded such genetic factors from our analyses, and the subsequent GEAs might thus largely underestimate the number of adaptive genes in our system. This might explain why we found so few genes associated with elevation, while altitude is well known to influence key traits such as plant size (Moles et al. 2009), flowering time (Kooyers et al. 2015; Vidigal et al. 2016; Wadgymar et al. 2018) or freezing tolerance (Zhen and Ungerer 2008). In addition, the confounding effect of population structure and adaptation at a regional scale may further mask the effect of the environment, as we recently showed for flowering time genes (Minadakis et al. 2023). As we also observed substructure within genetic clades (Fig. S3), within-genetic clades GEAs combined with common garden experiments might be more pertinent to identify population-specific loci. Furthermore, most traits exhibit a polygenic architecture, and detecting a large number of variants with subtle effects (for review Yeaman 2022) can be challenging with classical GEA or genome-wide association analyses (de Miguel et al. 2022). Hence, many crucial questions remain open to understand the genetic architecture as well as the geographical scale of adaptation in *B. distachyon*. With this diversity panel, the plant community is provided with a new means to unlock the source of natural variation in a monocot adapted to arid climate, beyond what previous pan-genome and QTL mapping studies permitted in this system.

## Materials and Methods

### Sampling and genotyping

Seeds from 299 *Brachypodium* plants were collected in 2018 and 2019 from southern France, Greece, and Montenegro. A representative subset of 110 plants was grown from seeds in greenhouse conditions, and DNA was extracted from their leaves with a DNeasy plant kit from Qiagen, following the manufacturer instructions. To confirm that only *B. distachyon* individuals were selected for whole genome sequencing, plants were genotyped using the microsatellite marker ALB165 (Giraldo et al. 2012) and by Sanger sequencing the GIGANTEA gene (López-Alvarez et al. 2012). Out of the 88 accessions identified as *B. distachyon*, 71 were selected for sequencing, while the remaining accessions were excluded from the rest of the analysis. The libraries were prepared and sequenced using Illumina HiSeq2500 (150 bp PE) by Novogene. Information about additional accessions collected and genotyped by genotyping-by-sequencing were obtained from Wilson et al. (2019) and displayed on a map with QGIS (version 3.4.13) together with all the *B. distachyon* used in this study.

### Sequencing, SNP calling, and filtering

Paired-end reads from the 71 newly sequenced accessions, together with the 65 *B. distachyon* accessions sequenced by Gordon et al. (2020), were aligned to version 3.0 of the *B. distachyon* Bd21 reference genome (https://phytozome-next.jgi.doe.gov) with bwa mem version 0.7.17-r1188 (Li 2013). Aligned reads were converted to bam files and sorted with samtools version 1.7 (Li et al. 2009), while accessions with multiple lanes were merged using the same program. Read duplicates were then removed with PICARD MarkDuplicates version 2.23.3, and variants were called with GATK v. 4.1.2.0 (McKenna et al. 2010). The resulting gvcf files were combined with the 196 gvcf files from the study of Stritt et al. (2022), leading to a dataset of 332 accessions. Subsequently, only SNPs with quality-by-depth higher than 8 and Phred-score more than 20 were kept using VCFTOOLS version 0.1.15 (Danecek et al. 2011).

### Population genetic structure and demographic history

The genetic structure of the samples was estimated by using principal component analysis and admixture analysis. To obtain independent SNPs, they were filtered for minor allele frequency > 0.05 and linkage disequilibrium < 0.4 using the function snpgdsLDpruning of the R package SNPRelate (Zheng et al. 2012). The principal components were calculated using the resulting 15,960 independent SNPs and the function snpgdsPCA of SNPRelate. On the same dataset, ancestry coefficients were estimated using the function sNMF of the R package LEA (Frichot and François 2015). Results for 2-10 K values for 20 replicates per K were retained, and the optimal K value was determined by calculating *ΔΚ* as described by (Evanno et al. 2005).

We obtained population size history using Relate v1.1.7 (Speidel et al. 2019) while filtering for FS > 60, SOR > 3, MQ > 40, -5.0 < MQRankSum < 5.0, QD < 2, ReadPosRankSum < -4.0, INFO/DP > 16,000. We also excluded genotypes with a genotype quality GQ < 20, an individual depth DP < 8x, and excluded sites with more than 20% missing genotypes, but importantly did not filter for a maf. This resulted in 6,728,435 SNPs. Relate also requires alleles to be polarized into ancestral and derived alleles. To do so, we used the genotypes from the ancestral C clade (Stritt et al. 2022), assuming the most frequent allele in that group to be ancestral. We then assigned each allele in the four other clades as ancestral or derived. We assume that individuals are strongly inbred, each owning two copies of the same haplotype. We therefore kept only one haplotype per individual. We used the Relate package (Speidel et al. 2019) to infer genome-wide genealogies and coalescence rates across all individuals. To take into account local variation in recombination rates, we used a recombination map for *B. distachyon* obtained from Huo et al. (2011). To obtain demographic estimates of times and population sizes, we used a mutation rate of 7x10-9 substitutions/generation (Lynch et al. 2016) and a generation time of one year as *B. distachyon* is annual. We set the prior for the effective haploid population size to 75,000 individuals based on previous estimates (Stritt et al. 2022). We subsequently fitted a time-varying population size history and inferred genome-wide topologies using the EstimatePopulationSize.sh script provided with the Relate package.

For each annotated gene (https://phytozome-next.jgi.doe.gov), the nucleotide diversity (pi) was calculated using pixy version 1.2.7.beta1 in 5 kb windows (Korunes and Samuk 2021). Tajima’s D was derived from these pi values in R using theta(pi) and the number of segregating sites.

### Estimating divergence times

We used the multispecies coalescent approach implemented in bpp v.4.2.9 (Rannala and Yang 2003; Flouri et al. 2018) to estimate the age of the splits among genetic lineages and clades, as also described in Stritt et al. (2022). 28 accessions (Cm4, Ren22, Msa27, Lb13, BdTR7a, 2_14_13, 4_52_6, 1c_25_14, Luc1, ABR6, Tso18, Bd30-1, BdTR2B, BdTR3C, 1a_32_12, Bd21, Mca12, Cro24, Cb23, San12, Arm-Arm-2B, Geo-G30i2, Geo-G31i4, Alb-AL1A, Ko2, MonSV13, Myt1, Vyt1) were selected to represent the five genetic clades. 200 random genomic regions of 1 kb length and at least 100 kb apart were chosen. For each of the 28 accessions, the 200 sequences were obtained by calling consensus sequences from the respective bam file. Inverse gamma priors were set to (3, 0.014) for the root age τ and to (3, 0.002) for the population size parameter θ, which corresponds to a mean theta of 0.001 and a mean root age of one million years, assuming a constant mutation rate of 7 × 10−9 substitutions per site per generation. The rooted species tree was defined as (((A_East, A_Italia), (B_East, B_West)), C_Italia), as inferred by Stritt et al. (2022). The MCMC was run four times independently, each time with 408,000 iterations, including a burn-in of 8,000 iterations. Highest posterior density intervals (HPDI) were estimated with the R package HDInterval version 0.2.4 (Meredith and Kruschke 2022). Relative cross-coalescence extracted from the output of Relate were also used to estimate divergence times, with relative cross-coalescence < 0.5 considered as populations being fully separated.

### Ecological niche modeling

Raster maps were downloaded for all bioclimatic variables from WorldClim 2.1 (Fick and Hijmans 2017) at a 30 sec resolution (about one kilometer). Global aridity index raster maps were obtained from https://cgiarcsi.community/data/global-aridity-and-pet-database/. We used the geolocations of the 332 accessions to extract bioclimatic variables at each locality with the R package raster 3.5-29. The 19 classical WorldClim variables (bio1 to bio19) and elevation were used unmodified. New variables were created from monthly values for precipitation, solar radiation, average temperature, maximum temperature, minimum temperature, and aridity, by calculating the averages of four months of putative biological interest (March to June, and November to February) (Table S1). Elevation data were downloaded using Google Earth. Correlations among bioclimatic variables were plotted as a bubble plot with the R package corrplot Version 0.92 (Wei and Simko 2021).

We used ecological niche models (ENMs) to estimate the climatic niches of the five genomic clades of *B. distachyon* and to investigate the degree of climatic niche overlap in both current and past conditions. As inputs for the models, we used the geographical coordinates of sequenced individuals and a set of environmental variables that were selected based both on biological relevance and on minimum collinearity between them: minimum temperature averaged from November to February, precipitation levels averaged from March to June, solar radiation levels averaged from March to June and elevation. The ENMs were produced with the MaxEnt software version 3.4.4 (Phillips et al. 2006), which uses machine learning to predict potential geographic distributions of species across different landscapes (Elith et al. 2006). The strength of association between the current distribution of different genomic clades and present climate variables was tested following the methods used by Williams et al. (2015) and Skalska et al. (2020). In brief, ENMs were considered to have identified relationships between the distribution of *B. distachyon* and the environmental predictors that were stronger than expected by chance given the spatial patterns in the data if the Area under the Curve (AUC) values of the real model were in the top five of the 100 models (99 null and 1 real), as described by Beale et al. (2008).

The ENMs were visualized using QGIS version 3.4.13-Madeira and maps were created to show the predicted suitable habitats for each clade. ENM modeling was restricted to the relevant study area by using a prepared mask (seen in Fig. 2) which was informed by very broad estimates of the extent of the sampling efforts (Fig. S1) and field observations. This included mainland Spain, Portugal, Turkey, Greece, Slovenia, Croatia, Bosnia, Montenegro, Albania, North Macedonia, Bulgaria, Georgia, Armenia and Azerbaijan. In France, *B. distachyon* is only found in the South (https://www.tela-botanica.org/bdtfx-nn-10075-synthese) and we only included the Occitanie and Provence-Alpes-Côte d’Azur administrative regions within the mask, rather than the whole country. For the same reason, we only included the Italian administrative regions to the south of Torino, Milan and the west of the Po valley. Sardinia, Corsica and the Balearic Islands were excluded. The dataset also included three sample points in Northern Iran and Iraq. No suitable administrative region could be found to effectively delineate these areas. We therefore created a buffer (1 decimal degree radians) around the sampling points.

To study past clade distributions, we projected current climatic niches onto paleoclimate maps using variables with a resolution of 2.5 arc-min for the Last Glacial Maximum (LGM) obtained from WorldClim1.4 (Fick and Hijmans 2017). The atmospheric general circulation models used in this study are the Community Climate System Model version 4 (CCSM4) (Gent et al. 2011). Past Maxent model projections were ‘clamped’ which restricts them to the range of climatic conditions that are seen in the current climate conditions (where the model was trained). This is recommended to avoid spurious model extrapolations when projecting into novel environments (Elith et al. 2010). Clamped areas can be seen coloured grey in Fig. S6.

The program ENMTools version 1.3 (Warren et al. 2010; Warren et al. 2021) was used to test whether there are statistically significant ecological differences between the clades, based on the ENMs produced by Maxent. Niche similarity was first calculated between each clade combination for the empirical data. 99 pseudoreplicate niche comparisons between each clade were then produced by pooled and randomized occurrence points of the empirical data. The overlap between ENMs generated from the empirical data for each clade comparison was then compared to the null distribution obtained using the pseudoreplicates, using Schoener’s D and the I statistic (Warren et al. 2010). We concluded that ENMs produced by two clades are significantly different if the empirical data had values smaller than the 95 of the pseudoreplicate values.

### Genotype-environment association studies

We used GEMMA 0.98.5 (Zhou and Stephens 2012) to fit a linear mixed model and test for association between SNPs and the 32 bioclimatic variables described above, while correcting for population structure. Α centered relatedness matrix was first produced with the option -gk 1. Association tests were performed using the option -maf 0.05 to exclude SNPs with minor allele frequency with values less than 0.05, as power is lacking for detecting associations using rare alleles (Marees et al. 2018).

We applied a False Discovery Rate (FDR, Benjamin and Hochberg 1995) threshold to control for the expected false positives rates among the rejected null hypotheses. To further reduce false positives, a sliding window approach was implemented across the whole genome, with window size of 5 kb and 2.5 kb overlap, using the R package rehh version 3.2.2 (Gautier et al. 2012). The selected windows contained at least two SNPs above the FDR threshold. We then used the BEDTOOLS version 2.26.0 intersect option (Quinlan and Hall 2010) to extract genes located in or overlapping with significant regions based on the v.3.2 annotation of the *B. distachyon* (https://phytozome-next.jgi.doe.gov).

### Scans of selection

The XTX analysis was performed with BayPass using the five genetic clades as focal populations (Gautier 2015). We generated the input file by using vcftools --count to calculate the allele frequency of each SNP present in our vcf (no filtering on minimum allele frequency). We then ran Baypass on our actual dataset with the following parameters: -pilotlength 500 -npilot 15 -burnin 2500 -nthreads 6. Top 0.1% outliers were chosen as a threshold of significance, as more conservative than the threshold calculated with a pseudo-observed dataset simulated with the co-variance matrix (Gautier 2015). We also used a sliding window approach, with window size of 5 kb and 2.5 kb overlap, using the R package rehh (Gautier et al. 2012).

### Age estimates and gene ontology enrichment

The age of each single SNP was computed with GEVA (Albers and McVean 2020) to estimate the average SNP age for each annotated gene (https://phytozome-next.jgi.doe.gov). All private SNPs to the combined A and B lineages were polarized using the ancestral C lineage using custom R scripts. GEVA was run on the five main scaffolds (corresponding to the five chromosomes) using the genetic map produced by (Huo et al. 2011) and the polarized SNP dataset. The average age of genes associated with the 32 bioclimatic variables described above or outliers in the XTX analysis was compared to the average age of genes at the genome wide level with Wilcoxon test. We only used the gene lists produced by the GEA and the XTX analyses if they contained at least fifteen genes. Gene Ontology search was performed with PANTHER v17.0 (http://www.pantherdb.org) using the built-in database for *B. distachyon*.

## Acknowledgements

We would like to thank Chris Thomas and Léa Frachon for valuable discussions on niche modelling and GEA respectively. We also thank Josep Casacuberta, Thibault Leroy and an anonymous reviewer for their time and comments on our manuscript.

## Author contribution

NM conceived the study, performed the niche modeling, population genetics and GEA analyses and wrote the manuscript. RH and YB performed age estimates, population size evolution and statistics. CS, DC and MT collected samples. HW performed the niche modeling analysis. ACR conceived the study and wrote the manuscript. All authors have read and agreed to the published version of the manuscript.

## Data, scripts, code, and supplementary information availability

Seeds will be distributed through GRIN but can in the meanwhile be obtained in small quantities upon request. Illumina paired-end sequencing data generated for this project are archived on the European Nucleotide Archive, project number PRJEB61986. Archive numbers for the reads produced by Gordon et al. (2018, 2020), Skalska et al. (2020) and Stritt et al. (2022) are available in the respective publications. Scripts are available at https://github.com/nminad/env_genomics.

