## Supplementary_figures for "The demographic history of the wild crop relative *Brachypodium distachyon* is shaped by distinct past and present ecological niches"

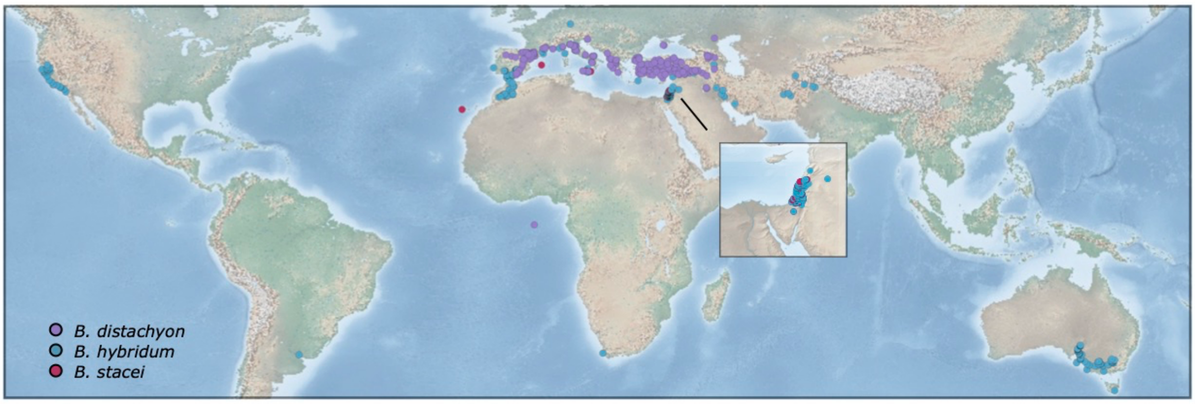


**Figure S1 - Geographic distribution of *B. distachyon*, *B. hybridum* and *B.stacei*.** The distribution is based on the 1897 samples genotyped by Wilson et al (2019) and the 332 samples described in this study.

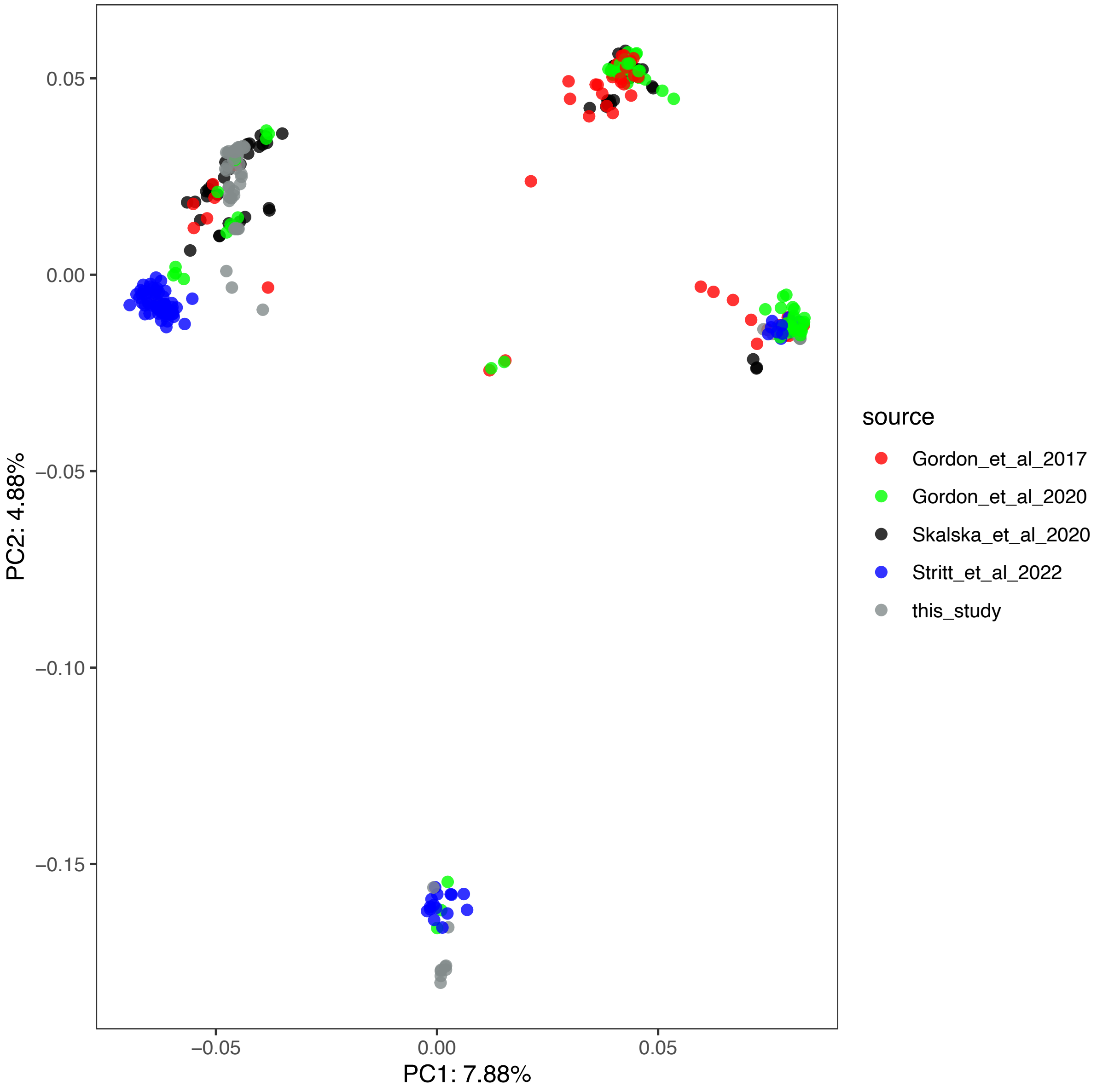


**Figure S2 –** PCA analysis performed with pruned SNPs for the 332 *B. distachyon* accessions described in the study (identical dataset as in Figure1). The colors indicate the study of origin.

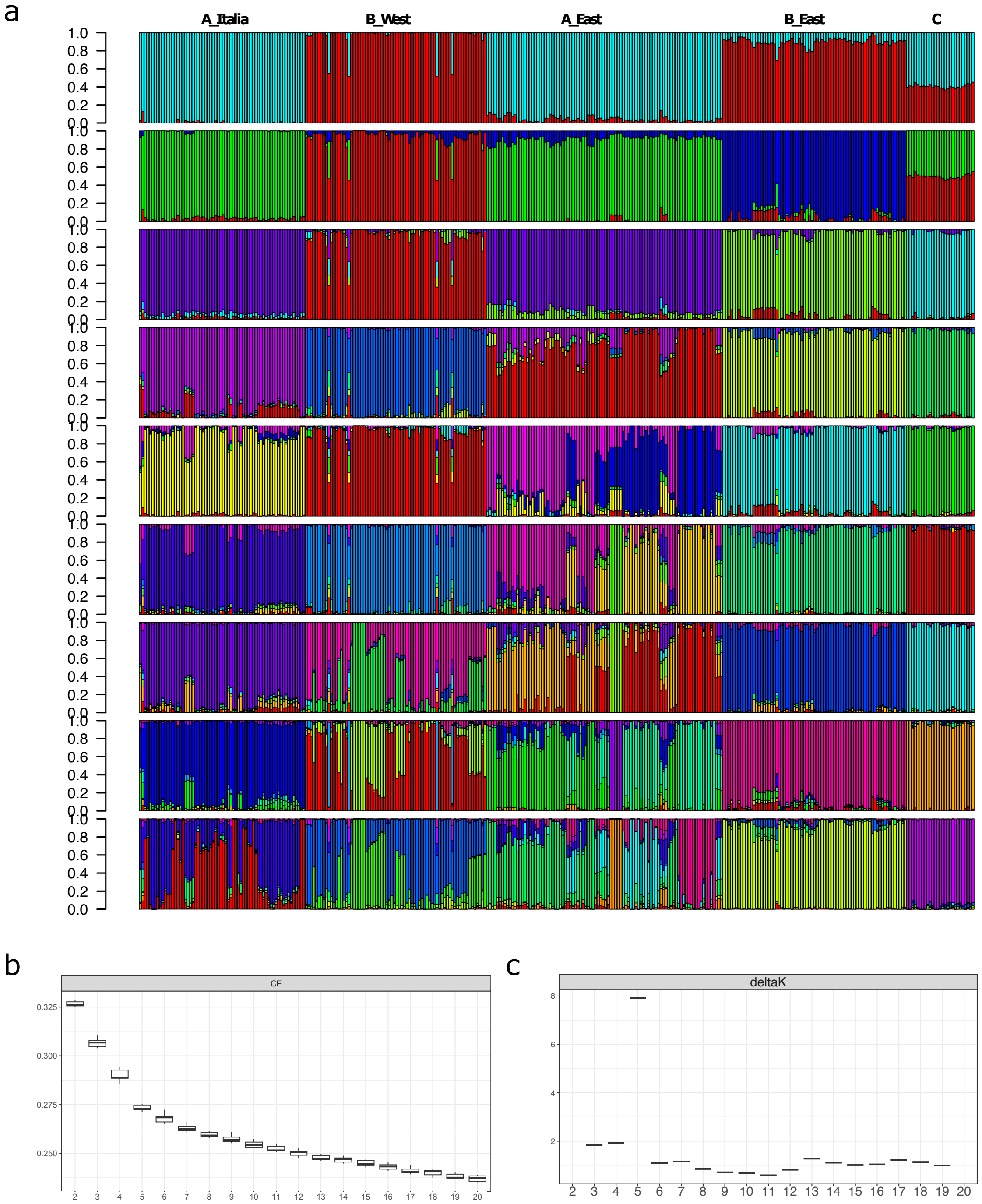


**Figure S3: sNMF analysis** a) sNMF analysis with K from 2 to 10 b) Cross-entropy with K from 2 to 20. c) Evaluation of the most conspicuous level of population structure. The calculations are performed as described in Evanno et al. 2005, with the difference that likelihoods are replaced by cross entropy (CE), the measure of model fit used by sNMF. Distributions for single values of K represent the 20 repetitions performed for each K. The low DeltaK values should not be interpreted as a low support, since the range of DeltaK values is expected to be different based on the cross-entropy values than those directly computed on the likelihood estimates.

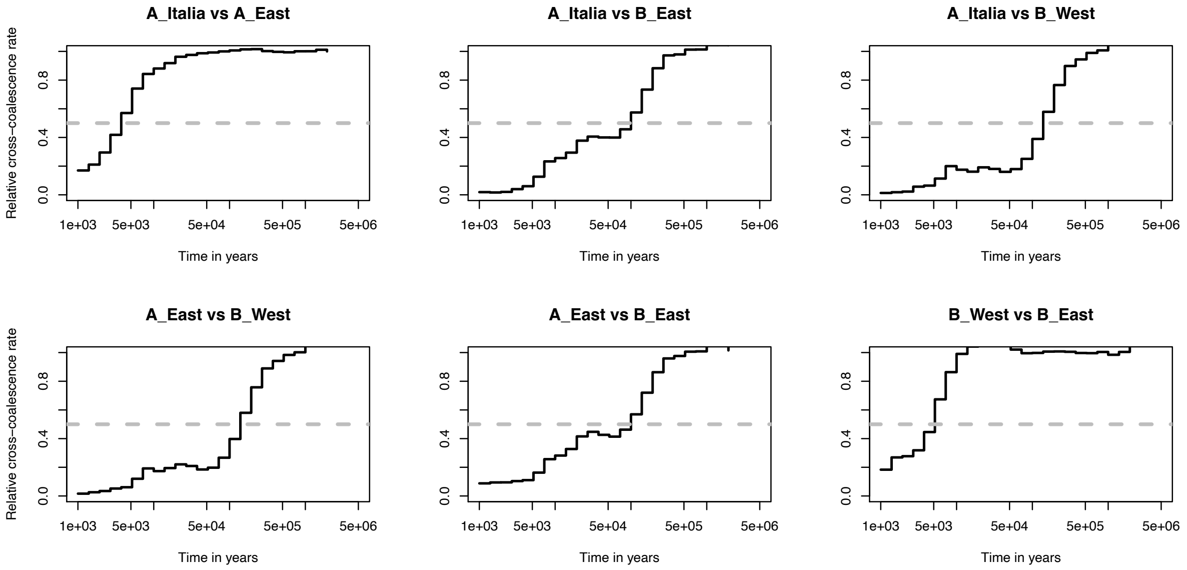


**Figure S4: Relative cross-coalescence rate between pairs of genetic clades.** The C clade was used to polarize the data and is not depicted here.

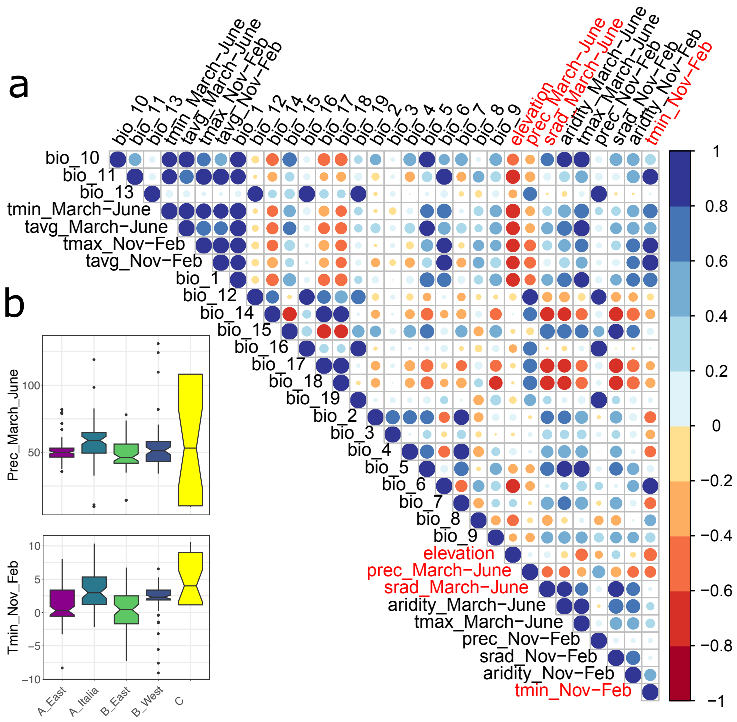


**Figure S5 - 32 Bioclimatic variable plots.** a) Correlation plots among 32 bioclimatic variables. Variables highlighted in red are the ones tested for niche modeling analyses b) Variation of prec_march_June and tmin_Nov_Feb across the four genetic clades.

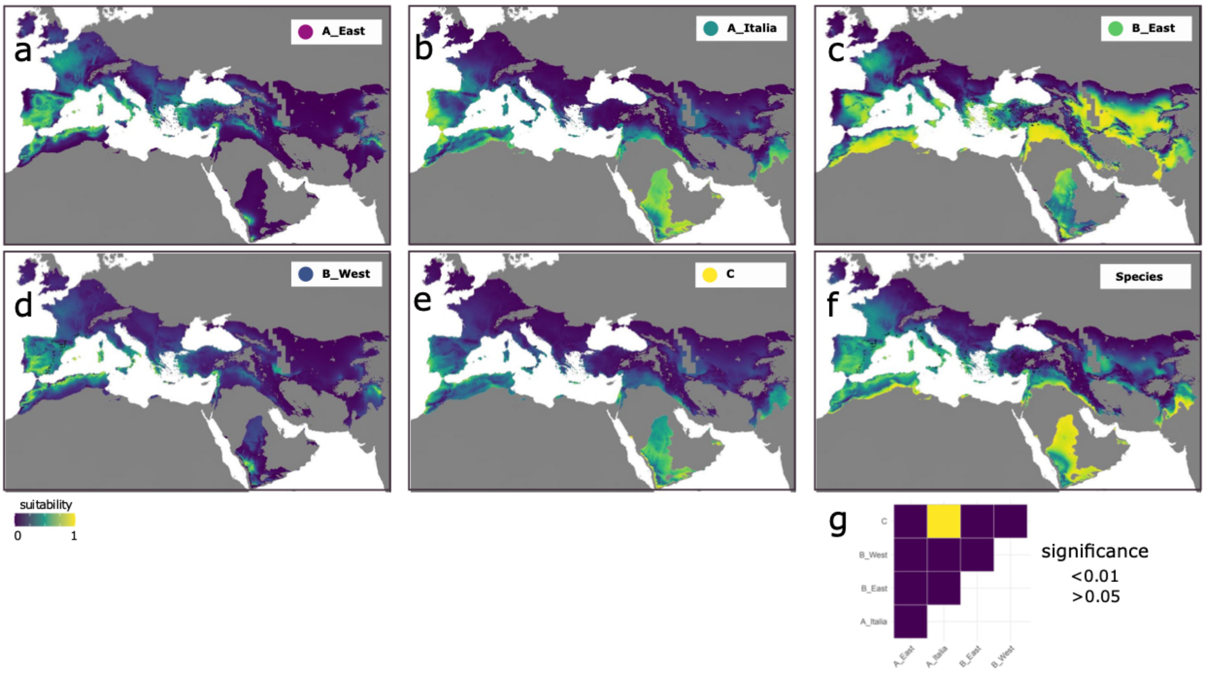


**Figure S6 - Niche modeling for last glacial maximum (LGM).** The maps display environmental suitability for the five genetic clades a) A_East b) A_Italia c) B_East d) B_West e) C and f) at the species level.

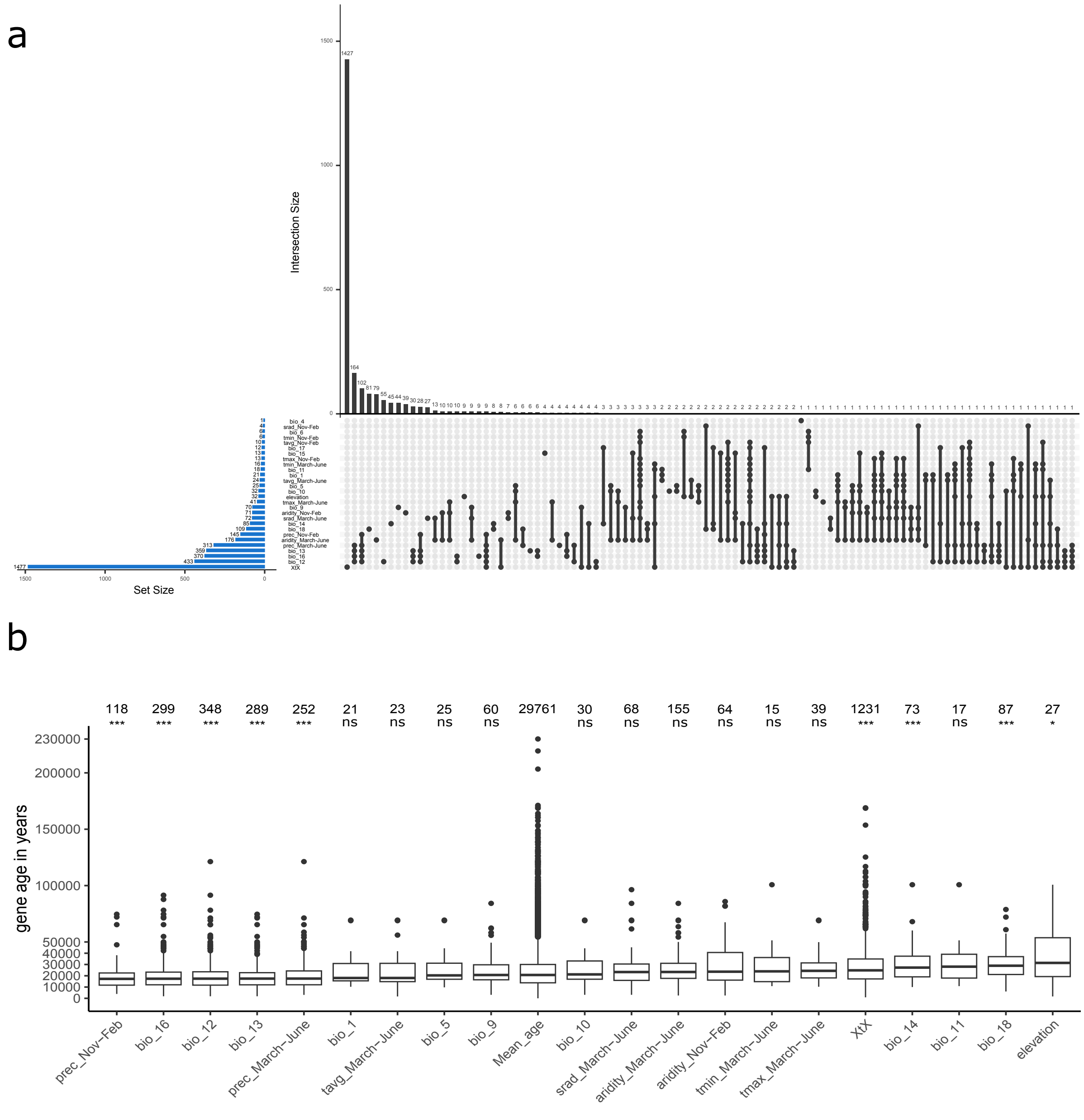


**Figure S7 - Genotype-environment associations analyses with all 32 bioclimatic variables.** a) The upset plot displays the gene set overlap among the 32 GEA and XtX analyses. b) Age estimates of genes associated with bioclimatic variables or under positive selection compared to the 29761 remaining annotated genes in the genome (total gene). The numbers at the top represent the number of genes analyzed per variable. Y-axis the variable ordered based on mean. * indicate whether the age of specific gene sets is significantly different than the age of the total gene set.

ns = no significant, * = p ≤ 0.05, ** = p ≤ 0.01, *** = p ≤ 0.001; Wilcoxon signed-rank test corrected for multiple testing.
